# Combinatorial small molecule-mediated activation of CBP/p300 with environmental enrichment in chronic severe experimental spinal cord injury to enable axon regeneration and sprouting for functional recovery

**DOI:** 10.1101/2021.06.07.447349

**Authors:** Franziska Mueller, Francesco De Virgiliis, Guiping Kong, Luming Zhou, Elisabeth Serger, Jessica Chadwick, Alexandros Sanchez-Vassopoulos, Akash Kumar Singh, Muthusamy Eswaramoorth, Tapas K Kundu, Simone Di Giovanni

## Abstract

The interruption of spinal circuitry following spinal cord injury disrupts neural activity and is followed by a failure to mount an effective regenerative gene expression response resulting in permanent neurological disability. Functional recovery requires the enhancement of axonal and synaptic plasticity of spared as well as injured fibres, which need to sprout and/or regenerate to form new connections. Here we will investigate whether the combined epigenetic stimulation of the regenerative gene expression program with neuronal activity-dependent enhancement of neuroplasticity and guidance can overcome the current inability to promote neurological recovery in severe and chronic spinal cord injury. This will be carried out by delivering the small molecule CBP/p300 activator CSP-TTK21 in combination with environmental enrichment. We will deliver the CBP/p300 activator CSP-TTK21 weekly between week twelve and twenty following a severe transection model of spinal cord injury in the mouse. Data analysis will assess modifications in regenerative signalling, in motor and sensory axon sprouting and regeneration, in synaptic plasticity as well as in neurological sensorimotor recovery. This approach will advance our mechanistic understanding of regenerative failure and provide insight of whether combining growth promoting strategies with enhancement of neural activity is an effective therapeutic strategy in chronic spinal cord injury.

## Introduction

Spinal cord injury (SCI) is a devastating disease affecting millions of people worldwide. Severe SCI leads to permanent motor, sensory, and autonomic dysfunction that disrupts the quality of life of affected people. The management of severe SCI is nowadays limited to supportive care. Current physical rehabilitation has measurable but limited benefits after moderate SCI but fails to improve recovery after more severe and chronic injuries. This permanent loss of function is primarily caused by disruption of the connectivity of long-distance and intraspinal axonal fibres that fail to regenerate and reconnect with the neural circuitry below the injury[1]. To date, the lack of axonal regeneration after injury has been attributed to two main interconnected factors: (i) the formation of a cellular inhibitory environment that promotes growth cone collapse and (ii) the lack of an intrinsic regenerative response[2]. Furthermore, abnormal activity in spinal circuits below the injury contributes to a progressive deterioration of sensorimotor function[3] that is likely intensified in patients with long-standing chronic SCI. Therefore, neuromodulation/rehabilitation and axonal regrowth strategies seek to promote activity-dependent neuroplasticity to improve sensorimotor recovery.

Accumulating evidence suggest that increasing neuronal activity not only contributes to axonal sprouting, but it also strengthens synaptic plasticity and stimulates targeted axonal regrowth, favouring re-connectivity and functional recovery[4, 5]. Modulation of axonal plasticity and growth of both motor and sensory fiber tracts, including of sensory circuits below the spinal lesion, can be carried out by specific neurorehabilitation schemes to enhance recovery after SCI[5]. However, this increase in regenerative growth and sprouting only partially promotes functional recovery, and it remains insufficient for re-connectivity and reestablishment of function in severe spinal injuries.

Attempts to stimulate the axonal regenerative response within the injured central nervous system (CNS) have been only partially successful through the manipulation of independent transcription factors or cofactors, including c-JUN, pCREB, SMAD1, MYC, HDAC5, KLF4, KLF7 and STAT3[6-15]. Accumulating evidence shows that epigenetic modifications can contribute to the transcription-dependent enhancement of the regeneration programme in sensory axons or the injured optic nerve[16-19]. Specifically, we found that the histone acetyltransferase (HAT) p300/CBP associated factor (PCAF) acetylates the promoters of several regeneration-associated genes (RAGs) driving their expression after sciatic nerve injury and that PCAF overexpression promotes sensory axon regeneration across the injured spinal cord[20]. We and others have also shown that the histone acetyltransferase p300 can enhance optic nerve regeneration, promoting the expression of selected RAGs[17], while inhibiting class I histone deacetylases or HDAC3, partially promotes axonal regeneration of sensory axons following spinal cord injury[21]. However, modulation of these targets that were identified from injury-dependent paradigms have not translated into significant neurological recovery. We recently found changes in neuronal activity by housing mice in an enriched environment (EE) (large cages with toys, tunnels, running wheels, and enriched bedding) or following specific chemogenetic modulation of neural activity, induced epigenetic modifications, enabling active transcription and regenerative growth. We next established that the CREB-binding protein (CBP) is the lysine acetyltransferase involved in this activity-dependent plasticity. Importantly, we showed that delivery of a small molecule specific activator of CBP/p300 named CSP-TTK21 promotes regenerative gene expression, axonal regeneration, plasticity and functional sensorimotor improvement following acute SCI in rodents[22]. TTK21 is a HAT activator conjugated to a glucose-derived carbon nanosphere (CSP) able to cross the blood brain barrier, cell and nuclear membranes with peak nuclear expression 3 days post i.p. injection[23]. It was previously shown to promote neurogenesis and ameliorate memory deficits in tauopathy model in mice through increased histone acetylation and expression of genes involved in synaptic plasticity[23, 24].

In addition, we have recently found that housing mice in an EE following a severe transection of the thoracic cord promotes significant sensorimotor recovery. These experiments suggest that housing animals in an EE represents an “enriched” form of neurorehabilitation. EE enhances neuronal activity promoting sprouting and growth of sensory as well as motor axons, along with increasing synaptic contacts (Supporting Figure 1).

**Supporting Figure 1.**
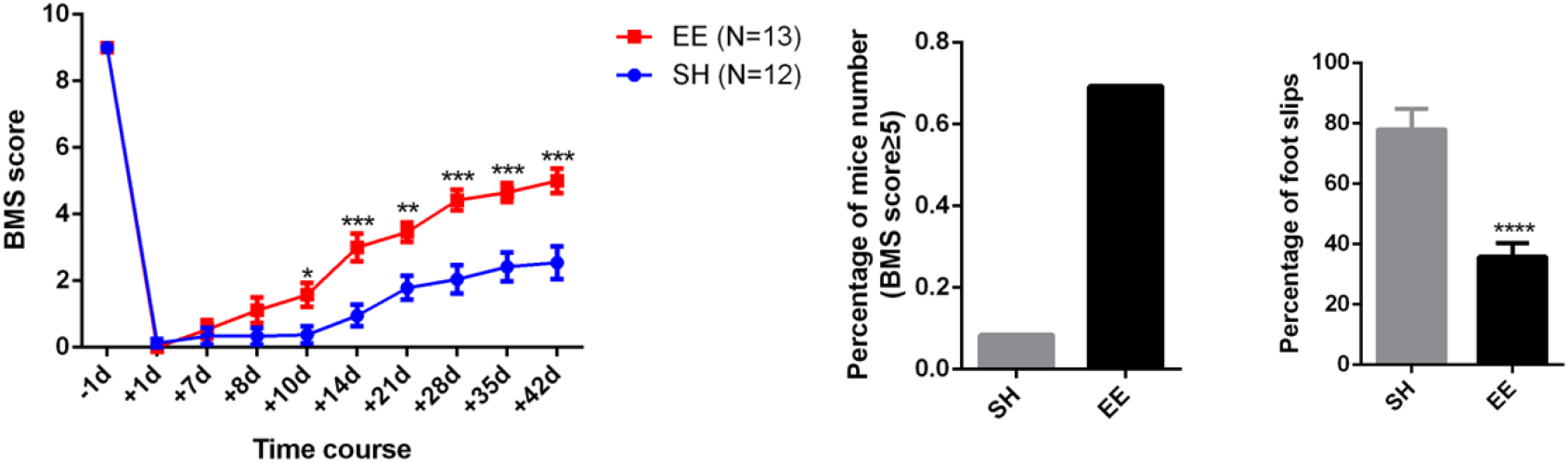
EE promotes functional neurological recovery after severe spinal cord transection. 12 weeks old mice were housed in standard housing (SH) or enriched environment (EE) one week after a spinal cord transection. Animals in SH remained impaired unable to step properly until day 42 after injury (BMS (left and middle graph) and Gridwalk (right graph). EE significantly enhanced locomotion (mean ± SEM, Two-way ANOVA, Fisher’s LSD post-hoc ** P<0.01; *** P<0.005; **** P<0.001).

However, the lack of integration between approaches aimed to promote regenerative molecular mechanisms with neurorehabilitation and neuromodulation after SCI remains a major limitation for repair in severe and chronic SCI, where reawakening a regenerative gene expression programme and stimulation of disrupted neural activity are especially challenging and potentially critical to repair and recovery. Additionally, mechanistic and therapeutic advances in chronic and severe spinal cord injury are especially rare and therefore represent a high priority.

## Hypothesis and relevance

Here we hypothesise that the pharmacological stimulation of CBP/p300 activity will enhance regenerative gene expression during a growth refractory phase twelve weeks after spinal injury, while housing animals in an EE one-week post-injury will stimulate neuronal activity, consolidate axonal and synaptic plasticity as well as promote directional regrowth. Since EE is believed to enhance activity in both descending motor axons and facilitate motor control through the recruitment of proprioceptive feedback circuits, which are the pathways targeted by CBP/p300 activator, we expect a potential strong synergy between the drug and EE.

Therefore, we postulate that the combination of the CBP/p300 activator CSP-TTK21 with EE following the most clinically relevant severe and chronic spinal cord injuries might enhance neuronal plasticity, regeneration and functional recovery, providing a better understanding of regenerative failure and filling a gap in the path to translation. In fact, the long-ranging implications of this work lie on testing a novel platform for combining pharmacology and neurorehabilitation to promote functional recovery after severe and chronic spinal cord injury, delivering the necessary pre-clinical evidence to support future clinical translation. Importantly, lack of validation of this hypothesis will also provide essential information allowing the scientific community to entertain alternative hypotheses. They include the possibility (i) that increases in regenerative gene expression and neural activity might not be synergistic in providing growth and plasticity; (ii) that recovery and repair might need task-specific neurorehabilitation; (iii) that re-establishment of functional circuitry needs targeting the spinal synapses; and finally (iv) that identification and manipulation of the extra-neuronal wound healing and scarring processes in chronic SCI might need to be engaged along with favouring neuroplasticity.

## Experimental design

The experimental design will ask the question of whether activation of CREB-binding protein (CBP) and the cognate protein p300 with the weekly systemic delivery of TTK21 combined with EE in chronic severe spinal cord transection injury in mice will allow enhancing gene expression that supports plasticity and regenerative growth, consolidate axonal and synaptic plasticity as well as promote directional regrowth. Since no data is available on TTK21 in chronic SCI at this stage, a severe transection model in mice has been chosen versus a contusion in rats because (i) it will allow a more accurate anatomical definition of axonal regeneration and sprouting; (ii) it has much less variability allowing for a more reliable statistical assessment keeping the number of animals relatively limited and (iii) it will need a fraction of the drug required (in mice vs rats) that is produced by our long-standing collaborator Professor Tapas Kundu (Centre for Drug Discovery, India). Currently the availability of the compound does not allow to initiate studies in rats in the short term, however these studies will surely prompt further investigation in severe chronic contusion in rats.

Specifically, adult 10-12 weeks old B6 mice will receive a spinal cord T9 severe transection injury that will destroy the ascending sensory fibres in the dorsal columns bilaterally as well as the most of the descending corticospinal, raphespinal, rubrospinal and reticulospinal motor tracts, mimicking severe clinical spinal cord injury. This lesion will lead to permanent severe impairment in sensorimotor function, severely limiting stepping up to at least 42 days post-injury, resembling clinical severe spinal cord injury (Supporting Figure 1). All mice will be placed in an EE one week after the spinal cord injury. The CBP activator TTK21 is coupled to slow-release carbon nanoparticles (CSP-TTK21) that allows a weekly administration maintaining stable activity levels in target tissues. Thus, CSP-TTK21 will be administered i.p. once per week starting twelve weeks after injury for eight to ten weeks. A control group of mice will be treated with nanoparticles and vehicle alone. The CSP-TTK21 and control nanoparticles will be used at the dosage of 20mg/kg that showed efficacy in subacute spinal cord injury as published recently[22].

Animals will be sacrificed at week 20-22 post-injury. Sprouting and regeneration of the dorsal columns will be analyzed with the retrograde axonal tracer Dextran-AF488 (injected in the sciatic nerve in proximity of L4-L6 DRGs 7 days before sacrificing the animals) that allows visualization of ascending fibres in the dorsal columns into and across the injury site as previously shown in Di Giovanni’s lab[22]. Sprouting and regeneration of the corticospinal tracts (CSTs) will be analysed with stereotaxic injections of the neural tracer BDA into the motor cortex 2 weeks before sacrifice, which allows visualization of descending fibres into and across the injury site according to standard procedures in Di Giovanni’s lab[25]. Axonal dieback as well as the number of fibres past the lesion site will be normalized to the number of labelled fibres prior to the lesion as previously shown[25]. Given it closely correlates to locomotion, we will also measure sprouting and regeneration of serotoninergic raphe-spinal motor tracts with 5-HT immunohistochemistry as recently described[22]. To assess whether the CSP-TTK21 treatment will enhance synaptic plasticity, we will measure the number of inhibitory VGAT or excitatory VGLUT1/2 synaptic terminals in proximity of neuronal targets such as interneurons in the dorsal horns and motoneurons in the ventral horns of the spinal cord (ChAT or NeuN immunostaining) including in association with specific tracing of CST, sensory or 5-HT fibres. This will be carried out by fluorescent multi-labelling experiments that will be analyzed by confocal microscopy. Histone acetylation as read out of CBP/p300 activation will also be evaluated in layer V neurons, raphe nuclei and DRG neurons by immunofluorescence. The expression of several regeneration associated genes including ATF3, JUN, GAP43, SPRR1a, KLF family members, and STAT3 will also be studied by immunofluorescence in sensory and motor neurons.

Lastly, we will assess locomotion, coordination and sensorimotor integration by performing open field assessment with the BMS and the gridwalk tests, as recently shown[22]. In addition, Von Frey test for mechanoception and mechanical allodynia as well as Hargreaves test for thermoception and thermal hyperalgesia will be used to specifically assess the function of the ascending sensory tracts.

## Summary Table

| Research question | Hypothesis | Sampling plan | Statistical analysis | Outcomes that confirm or disconfirm hypothesis |
| --- | --- | --- | --- | --- |
| Does weekly systematic CSP-TTK21 delivery in combination with EE after a chronic severe spinal cord injury promote *sensorimotor fiber regeneration*? | We hypothesize that combinatorial treatment of CSP-TTK21 and EE will enhance sensorimotor fiber regeneration due to the synergy between the two treatments enabling fibres to regenerate through the site of injury and provide directional regrowth.<br>Since EE is believed to enhance activity in both descending motor axons and facilitate motor control through the recruitment of proprioceptive feedback circuits, which are the pathways targeted by CBP/p300 activator, we expect a strong synergy between the drug and EE. | TTK21 vs control: increase of the regeneration index at 300 μm beyond the lesion site from 20 to 40; SD= 8; Power 90%; P: 0.01, N: 8 or P:0.05, N: 6 | 5-HT Immunostaining: Normally distributed data will be evaluated using a two-tailed unpaired Student's *t*-test or a one-way ANOVA when experiments contained more than two groups. The Tukey's post-hoc test will be applied when appropriate. The Mann-Whitney's U-test will be used for non-parametric evaluation.<br>Quantification of axonal regeneration: Normally distributed data will be evaluated using a two-tailed unpaired Student's *t*-test or a one-way ANOVA when experiments contained more than two groups. The Tukey's post-hoc test will be applied when appropriate. The Mann-Whitney's U-test will be used for non-parametric evaluation.<br>A threshold level of significance $\alpha$ was set at P-value <0.05. Significance levels will be defined as follows: * P-value <0.05; ** P-value <0.01; *** P-value <0.001, **** P-value <0.0001. All data analysis will be performed blind to the experimental group. | Hypothesis would be strongly confirmed if significant regeneration of sensory and motor (CST, raphespinal axons) fibres is observed following CSP-TTK21 and EE treatment compared to control.<br>Additional evidence to support this hypothesis would be provided by showing enhanced synaptic plasticity via an increased number of inhibitory VGAT or excitatory VGLUT1 synaptic terminals in proximity of neuronal targets in the spinal cord. |
| Does the combination of CSP-TTK21 systemic weekly delivery and EE following a chronic and severe spinal cord injury enhance *functional recovery* | We hypothesise that combinatorial treatment of CSP-TTK21 and EE will promote functional recovery. | TTK21 vs control: changes in score from $\geq 1.5$; SD: 4 and 3; Power 90%; P:0.05; N= 10 | Gridwalk and BMS: Data will be evaluated using a repeated measures ANOVA with a Bonferroni or Sidak multiple comparison test.<br>A threshold level of significance $\alpha$ was set at P-value <0.05. Significance levels will be defined as follows: * P-value <0.05; ** P-value | Hypothesis would be strongly confirmed if significant difference was observed in BMS and gridwalk scores between CSP-TRTK21 and control. Particularly if a significant improvement of BMS score and reduction in the number of footslips was observed |
|  |  |  | <0.01; *** P-value<br><0.001, **** P-value<br><0.0001. All data analysis<br>will be performed blind to<br>the experimental group. | in the CSP-TTK21<br>group vs control<br>following weekly<br>injections. |
| Does the<br>combination of CSP-TTK21 systemic<br>weekly delivery and<br>EE following a<br>chronic and severe<br>spinal cord injury<br>promote *gene*<br>*expression and*<br>*histone acetylation*<br>to enhance<br>plasticity and<br>regenerative<br>growth | We hypothesise that<br>stimulating CBP/p300<br>activity will enhance<br>regenerative gene<br>expression and<br>histone acetylation<br>during a refractory<br>phase 12 weeks post<br>injury, while EE one<br>week post injury<br>stimulates neuronal<br>activity and<br>consolidates axonal<br>and synaptic plasticity. | TTK21 vs control:<br>change in<br>fluorescence<br>intensity of 1.5<br>fold; SD=0.25;<br>P:0.05; Power<br>90%; N=4 | IHC/IF: Normally<br>distributed data will be<br>evaluated using a two-tailed unpaired Student's<br>t-test or a one-way<br>ANOVA when<br>experiments contained<br>more than two groups.<br>The Tukey's post-hoc test<br>will be applied when<br>appropriate. The Mann-Whitney's U-test will be<br>used for non-parametric<br>evaluation.<br>A threshold level of<br>significance $\alpha$ was set at<br>P-value <0.05.<br>Significance levels will be<br>defined as follows: * P-value <0.05; ** P-value<br><0.01; *** P-value<br><0.001, **** P-value<br><0.0001. All data analysis<br>will be performed blind to<br>the experimental group. | Hypothesis would be<br>strongly confirmed if a<br>significant increase in<br>gene expression of<br>regeneration<br>associated genes (e.g.<br>c-Jun, Atf3, pStat3) as<br>well as increased<br>histone acetylation<br>(e.g. H3k27ac, H3k9ac,<br>H4k8ac) was observed<br>following CSP-TTK21<br>treatment versus<br>control. |

## Materials and Methods

### Mice

Mouse experimentation will be carried out in accordance with regulations of the UK Home Office under the Animals (Scientific Procedures) Act 1986, with Local Ethical Review by the Imperial College London Animal Welfare and Ethical Review Body Standing Committee (AWERB). C57Bl6 (Harlan, UK) mice ranging from 10 to 12 weeks of age will be used for all experiments. For all surgeries, mice will be anesthetized with isoflurane (4% induction 2% maintenance) and buprenorphine (0.1 mg/kg) and carprofen (5 mg/kg) will be administered peri-operatively as analgesic.

Animals will be kept on a 12 hour light/dark cycle with food and water provided ad libitum, at a constant room temperature and humidity (21° C and 5%, respectively). Standard housing for mice consists of 26×12×18 cm^3^ cages housing 4 mice with tissue paper for bedding, a tunnel and a wooden chew stick. The enriched environment housing consists of 36⨯18⨯25 cm^3^ cages housing 5 mice with tissue paper for bedding, a tunnel and a wooden chew stick. EE cages also host: additional nesting material which included nestlets, rodent roll and sizzle pet (LBS biotech); a hanging plastic tunnel (LBS biotech) and a plastic igloo combined with a fast-track running wheel (LBS-biotech); a wooden object (cube, labyrinth, tunnel, corner 15) (LBS biotech) that is changed every 5 days to help maintain a novel environment; and 15 g fruity gems (LBS biotech) every 5 days to encourage exploratory and natural foraging behaviour.

### Spinal cord injury (SCI)

Surgeries will be performed as we previously reported[25]. A laminectomy at vertebra T9 will be performed to expose spinal level T9 and a deep dorsal over-hemisection past the central canal leaving only sparing of the ventral white matter will be performed using micro-scissors (Fine Science Tools). For the sham surgery a laminectomy will performed but the dorsal over-hemisection will not. Between week twenty and twenty-two after spinal cord injury animals will be deeply anesthetized and perfused transcardially.

### CSP-TTK21 administration

Animals will be randomised to treatment with the CBP/p300 activator bound to carbon nanospheres (CSP-TTK21) or a control of just carbon nanospheres (CSP) (mice – 20 mg/kg injected i.p. once a week). Mice will receive the first IP injection 12 weeks after SCI until sacrifice.

### Neuronal tracing

For dorsal column tracing of sensory DRG axons, 2 µl of Dextran-AF488 (Life Technology) will be injected into the sciatic nerve bilaterally one week before sacrifice using the fire polished glass capillary which is connected with a 10 µl Hamilton syringe.

For anterograde tracing of motor corticospinal tracts, two weeks before sacrifice, mice that received a spinal cord injury will be injected with the axonal tracer BDA with a Hamilton micro-syringe in the motor cortex following standard stereotaxic coordinates as we previously described[26].

### 5-HT immunostaining

For 5-HT immunostaining we will follow a protocol we previously described [27]. Briefly, spinal cord sections will be post-fixed in 0.1% glutaraldehyde in 4% PFA and will next be quenched with ethanol peroxide and transferred to 0.2% sodium borohydride. They will undergo antigen retrieval in 0.1 M citrate buffer, pH 4.5, and microwave irradiation at 98 °C for 2 minutes. The sections will be incubated with rabbit anti-5-HT (1:8000, ImmunoStar) in 4% NGS in 0.3% TBS-Triton X100 for 4 days at 4 °C. Next, the sections will be incubated with an Alexa fluorescent secondary antibody. Finally, the sections will be coverslipped.

### Behavioral analysis

Mice will be trained daily for 2 weeks pre-injury before baseline measurements and then assessed on day 7 post-injury and weekly thereafter. All behavioral testing and analysis will be done by two observers blinded to the experimental groups.

#### Gridwalk

Mice will cross a 1 m long horizontal grid 3 times. Videos of the runs will be analyzed at a later time-point and errors from both hind-limbs will be counted. Error values represent the total number of slips made by both hindlimbs over the 3 runs.

#### Open-field test

The Basso Mouse Scale (BMS)[28] will be used to assess open-field locomotion. Each animal will be allowed to freely move in the open field for 4 min while two independent investigators blinded to experimental group will score it. The BMS score and sub-score will be given. Only the animals showing frequent or consistent plantar stepping in the open field (BMS score ≥5) will be tested on the grid walk.

#### Von Frey

The Von Frey test will determine the mechanical force required to elicit a paw withdrawal response. Each animal will be tested in each paw 3 times. Only animals showing plantar placement in the open field (BMS score ≥3) will be tested.

#### Hargreaves

The Hargreaves test will determine the latency of a thermal nociceptive stimulus required to elicit a paw withdrawal response; each animal will undergo 2 test sessions in which each paw will be tested 3 times. Only animals showing plantar replacement in the open field (BMS score ≥3) will be tested.

### Quantification of axonal regeneration

For each spinal cord after injury, the number of fibres rostral and caudal to the lesion and their distance from the lesion epicentre (depending on whether sensory or motor axons) will be analysed in 4–6 sections per animal with a fluorescence Axioplan 2 (Zeiss) microscope and with the software Stereo-Investigator 7 (MBF Bioscience). The lesion epicentre will be identified by GFAP staining in each section at 20× magnification. The total number of labelled axons or signal intensity of the traced axons rostral to the lesion site will be normalized to the total number of labelled axons or of the signal intensity caudal or rostral to the lesion site counted in all the analysed sections for each animal, obtaining an inter-animal comparable ratio. Sprouts and regrowing fibres will be defined following the anatomical criteria reported by Steward and colleagues[29].

### Histology and immunohistochemistry

Tissue will be post-fixed in 4% paraformaldehyde (PFA) (Sigma) and transferred to 30% sucrose (Sigma) for 5 days for cryoprotection, the tissue will then be embedded in OCT compound (Tissue-Tek) and frozen at -80°C. DRG, spinal cord and cortexes will be sectioned at 10, 20 and 20 µm thickness respectively using a cryostat (Leica). Immunohistochemistry on tissue sections will be performed according to standard procedures. For selected antibodies used, antigen retrieval will be performed submerging the tissue sections in 0.1 M citrate buffer (pH 6.2) at 98°C for 5 minutes. Next, tissue sections will be washed with PBS to remove the excess of citrate buffer and blocked for 1h with 8% BSA, 1% PBS-TX100. Finally, the sections will be incubated with anti-p-STAT3 (1:200, Rabbit, Cell Signaling Technology 9145), c-Jun (1:50, CST, #9165), ATF3 (1:100, Santa Cruz, sc-188), pErk (1:250, CST, #9101), IGF1R (1:200, CST, #3027), CTB (1:1000, List biological 703), Tuj1 (1:1000, Promega G7121), H4K8ac (1:1000, ab15823), H3K27ac (1:500, ab4729), H3K9ac (1:500, Cell Signalling 9671), GFAP (1:500, Millipore AB5804), CD68 (1:500, Abcam ab213363), vGlut1 (1:1000, Synaptic system 135302), vGAT1 (1:500, Synaptic systems 131011), antibodies at 4°C O/N. This will be followed by incubation with Alexa Fluor conjugated secondary antibodies according to standard protocol (1:1000, Invitrogen). Slides will be counterstained with DAPI to visualise nuclei whenever necessary (1:5000, Molecular Probes).

### Image analysis for IHC

All analysis will be performed by the same experimenter who was blinded to the experimental groups. Photomicrographs will be taken with a Nikon Eclipse TE2000 microscope with an optiMOS scMOS camera using 10x or 20x magnification using ImageJ (Fiji 64bits 1.52p), Macromanager 2.0 software for image acquisition or at 20× magnification with an Axioplan 2 (Zeiss) microscope and processed with the software AxioVision (Zeiss).

### Analysis of GFAP and CD68 intensity around the lesion site

GFAP and CD68 intensity and area with positive signal will be quantified from sagittal spinal cord sections from one series of tissue for each animal. Quantification will be done using ImageJ, the background will be subtracted and then the mean pixel intensity and area of immunoreactivity will be measured.

### Analysis of fluorescence intensity

For quantitative analysis of pixel intensity, the nucleus or soma of DRG or layer V cortical neurons will be manually outlined in images from one series of stained tissue for each mouse. To minimize variability between images pixel intensity will be normalized to an unstained area and the exposure time and microscope setting will be fixed throughout the acquisition.

### Analysis of 5HT fibres in the ventral horn

Intensity of 5HT immunohistochemistry will be measured in the ventral horn of L1-4 spinal sections. Quantification will be done using ImageJ, the background will be subtracted and then the mean pixel intensity will be measured from one series of tissue for each animal.

### Analysis of vGlut1 and VGAT immunohistochemistry in proximity to motor neurons

VGlut1 and VGAT synaptic boutons will be imaged with a SP8 Leica confocal microscope. Z-stacks images were taken with an average thickness of 15µm with a step-size of 0.3µm. Sequential line scanning was performed when more than two channels were acquired. Multi-fluorescent orthogonal 3D image analysis and visualization will be performed using Leica LAS X software. The average number of vGlut1 or VGAT boutons opposed to motor neurons in the ventral horn of L1-3 spinal sections will be calculated by analysing 20 motor neurons per replicate. All analyses will be performed blind to the experimental group.

### Statistical analysis

#### Measures for avoidance of bias (e.g. blinding, randomisation)

We adhere to the principles of NC3Rs and adhere to the ARRIVE guidelines.[30] All experiments will be performed in blind to the treatment and experimental group. Behavioural studies will be assessed by 2 independent investigators (e.g.: research associate and research assistant) blind to one another, to the treatment and to injury group when relevant. Randomization will follow a computerized sequence.

#### Sample size calculation, power calculations and justification of effect size

the size of our in vivo experimental groups as planned in our aims have been defined following the Animal Experimentation Sample Size Calculator (AESSC) tool. Sample size calculations were performed using two-tailed t-test and one-way ANOVA (significance ≤ 0.05; power ≥ 0.90; G*Power); N indicates number/group. The specific effect size has been estimated based upon similar studies showing significant differences between experimental and control group. Two animals will be added to each experiment based upon the probability of losing animals due to the experimental procedure such as spinal surgery. While selected N is derived by our power calculation it is also compatible and comparable with what has been shown in the literature for similar experiments. Exclusion criteria include tissues with low quality for further experimentation or imaging as well as animals with injury severity beyond 2 SD from the mean.

Specific calculations are found here below:

Immunofluorescence (TTK21 vs control): change in fluorescence intensity of 1.5 fold; SD=0.25; P:0.05; Power 90%; N=4

Axonal regeneration (TTK21 vs control): increase of the regeneration index at 300 µm beyond the lesion site from 20 to 40; SD= 8; Power 90%; P: 0.01, N: 8 or P:0.05, N: 6

Behavioral tests (TTK21 vs control): changes in score from ≥1.5; SD: 4 and 3; Power 90%; P:0.05; N= 10

Results will be expressed as mean ± SEM. Statistical analysis will be carried out using GraphPad Prism 9. Normality will be tested for using the Shapiro-Wilk test. Normally distributed data will be evaluated using a two-tailed unpaired Student’s *t*-test or a one-way ANOVA when experiments contained more than two groups. The Tukey’s post-hoc test will be applied when appropriate. The Mann-Whitney’s U-test will be used for non-parametric evaluation. A threshold level of significance α was set at P-value *<*0.05. Significance levels will be defined as follows: * P-value *<*0.05; ** P-value *<*0.01; *** P-value *<*0.001, **** P-value <0.0001. All data analysis will be performed blind to the experimental group.

